# Acute and sub-chronic toxicity of 6PPD-quinone to early-life stage lake trout (*Salvelinus namaycush*)

**DOI:** 10.1101/2024.03.26.586843

**Authors:** Catherine Roberts, Junyi Lin, Evan Kohlman, Niteesh Jain, Mawuli Amekor, Alper James Alcaraz, Natacha Hogan, Markus Hecker, Markus Brinkmann

## Abstract

N-(1,3-Dimethylbutyl)-N’-phenyl-p-phenylenediamine-quinone (6PPD-q) is a rubber-tire derivative which leaches into surface waters from roadway runoff, from tire particles and has been identified as a possible driver of urban runoff mortality syndrome in coho salmon. Sensitivity to this toxicant is highly variable across fish species and life stages. With environmental concentrations meeting or exceeding toxicity thresholds in sensitive fishes, the potential for ecologically relevant effects is significant. There is currently no data regarding the sensitivity of lake trout (*Salvelinus namaycush*) to 6PPD-q. As early-life stages of fishes are typically more sensitive than adults, the goal of these studies was to evaluate the acute and sub-chronic toxicity of 6PPD-q to early-life stage lake trout. Alevins exposed from hatch until 45 days post hatch (dph) to time-weighted average 6PPD-q concentrations ranging from 0.22-13.5 μg/L exhibited a 45-day median lethal dose (LC_50_) of 0.39 μg/L. Deformities throughout growth were observed, with a unique pooling of blood observed in the caudal fin and eye. A subsequent acute study with exogenously feeding lake trout fry determined a 96-hr LC_50_ of 0.50 μg/L. From these studies we can conclude that lake trout alevins and exogenously feeding fry are sensitive to 6PPD-q, which underscores the relevance of this chemical to inland freshwater ecosystems.

## INTRODUCTION

The emerging contaminant of concern, 6PPD-q (N-(1,3-Dimethylbutyl)-N’-phenyl-p-phenylenediamine-quinone), has become a recent focus of ecological concern, given its acute toxicity to several salmonid species at concentrations lower than what is found in urban stormwater runoff and receiving streams. In fact, it was recently discovered that 6PPD-q was responsible for pronounced pre-spawn mortality coho salmon (*Oncorhynchus kisutch*) within the Pacific Northwest of the United States have faced for decades, a trend linked to both roadway density and precipitation events ^1,2^. This phenomenon was termed Urban Runoff Mortality Syndrome (URMS) ^3^. A follow up study by Tian et al. ^4^ reported a 24-hour median lethal concentration (LC_50_) of 95 ng/L for 6PPD-q in coho salmon. 6PPD-q is a product of the environmental oxidation of N-(1,3-Dimethylbutyl)-N’-phenyl-p-phenylenediamine (6PPD), an antioxidant used in rubber, notably in tire tread. Consequently, 6PPD-q can enter waterways during precipitation events as tire wear particles (TWP) from roads are washed into aquatic environments. Studies in a number of urban settings have revealed environmental concentrations of 6PPD-q ranging from 0.08 to 2.85 μg/L ^5–7^. These concentrations surpass the LC_50_ for coho salmon and several other sensitive species studied so far, including rainbow trout (*Oncorhynchus mykiss*) (72-hour LC_50_ 1.00 μg/L), brook trout (*Salvelinus fontinalis*) (24-hour LC_50_ 0.59 μg/L) ^8^, and white spotted char (*Salvelinus leucomaenis pluvius*) (24-hour LC_50_ 0.51 μg/L) ^9^. Consequently, there exists a significant acute risk to wild populations of certain salmonid species; however, few of the globally known 66 distinct species of salmonids have been tested to date limiting current risk assessment efforts.

The broad range of sensitivity among fishes to this compound is noteworthy. To date, the only fish species found to be sensitive are in the Salmonidae family. However, while some salmonid species, such as white-spotted char and brook trout, exhibit sensitivity at very low concentrations of 6PPD-q in the sub-μg/L range, other closely related species such as Arctic char (*Salvelinus alpinus*) and southern Asian dolly varden (*Salvelinus curilus*) show no acute adverse effects even at very high non-environmentally relevant concentrations ^8,9^. At the same time, experimental data on acute lethality following exposure to 6PPD-q is lacking for most salmonid (and non-salmonid) species, and presently, there are no reliable predictors that allow risk assessors to identify species that are vulnerable to the exposure with this chemical. In addition, little is known in regard to early-life stage or chronic exposure of salmonid species to 6PPD-q ^10,11^. Given that embryonic and larval life stages of fishes often tend to be more sensitive to contaminants than adults ^12^, there is the potential that these life stages might be more susceptible to the lethal or sub-lethal effects compared to their adult conspecifics. Given that 6PPD-q can leach from TWPs, the risk exists that salmonid alevins, which are closely associated with gravel and sediment, may be of particular risk to exposure to 6PPD-q leaching from settled TWP. In addition, while aqueous exposure to 6PPD-q through urban stormwater runoff normally occurs in pulses ^7^, one might speculate that exposure of salmonids alevins through TWPs in sediments could occur over prolonged periods of time.

Lake trout (*Salvelinus namaycush*) are a freshwater species which is prevalent across North America, and typically found in cooler, deeper waters ^13^. Lake trout are long-lived, with lifespans surpassing 20 years ^13^. Despite their longevity, the status of lake trout populations in the Laurentian Great Lakes and across Ontario are of concern, with declining numbers and distribution of the species ^14^. While this decline is largely attributed to the adverse effects of climate change and habitat loss ^14^, there is the potential that lake trout populations could be subject to 6PPD-q via urban stormwater runoff. The Canadian Great Lakes basins, where lake trout reside, are subject to as high 2.23 billion m^3^ of total runoff from the Greater Toronto Area annually. While 6PPD-q specifically has not been studied in the Great Lakes, the Don River, which feeds into Lake Ontario, has been documented to contain up to 2.85 μg/L of 6PPD-q (Johannessen et al., 2021). If lake trout are sensitive to 6PPD-q exposure, particularly during their early-life stages, it could represent an additional threat to population recovery and maintenance efforts in these ecosystems. Therefore, lake trout are a crucial native species of concern in Canada and the United States.

In this study, we aimed to determine the sub-chronic and acute sensitivity of lake trout alevins and exogenously feeding fry, respectively, to exposure with 6PPD-q. In addition, this study observed and described a number of sub-lethal effects of 6PPD-q exposure, which can provide important insights into the mechanisms through which 6PPD-q exerts its high level of toxicity. Together, this study provides important new information for environmental risk assessors and regulators to understand the impacts of 6PPD-q on an important fish species with ecological, commercial, and cultural relevance in North America.

## MATERIALS AND METHODS

### Chemical Source

6PPD-q was sourced from Toronto Research Chemicals, Cat# P348790, Lot # 6-ABK-122-2; 97% purity. Powder stock was made up in dimethyl sulfoxide (DMSO), at 10,000x the exposure concentrations, resulting in a 0.01% DMSO (v/v) concentration for all treatments. Deuterium-labeled 6PPD-q d5 (Cat# P348691) was also obtained from Toronto Research Chemicals, and the neat compound was dissolved in LC-MS-grade methanol at 1 mg/L. Exposure solutions were freshly prepared daily.

### Lake trout source and housing

Lake trout eggs were sourced from fish collected from Whiteswan Lake (SK, Canada) in conjunction with the Fisheries Department of the Government of Saskatchewan, Canada. Eggs from ten different females were each fertilized using milt from two different males, resulting in 20 different genetic lines. Embryos were maintained in the dark at the University of Saskatchewan Aquatic Toxicology Research Facility (ATRF) at 10 ± 0.5°C in a flow-through heath tray system until they reached the eyed stage, checked twice daily for temperature and mortalities, which were removed, as well as water quality (DO, ammonia, NH_3_, NH_4_, hardness) (La Motte). Prior to hatching, embryos were transferred to larger glass aquaria for monitoring and randomized exposure tank allocation. All experiments involving animals were reviewed and approved by the University of Saskatchewan Animal Research Ethics Board (Animal Use Protocol 2022-0002). Lake trout embryos were transported to the University of Saskatchewan under the Government of Saskatchewan Fish Transport/Stocking permit FTP02-2022.

### Sub-chronic exposure

Tanks (2.5 L) were individually aerated and randomized on shelves and allowed to equilibrate with exposure water for 24 hours prior to exposure. Upon hatch, 15 lake trout alevins were placed randomly in each tank and were exposed to nominal concentrations of 6PPD-q of 0.625, 1.25, 2.5, 5, 10, and 20 μg/L, as well as a solvent control (0.01% DMSO) with five replicate tanks per treatment. Tanks were maintained under semi-static conditions, with a 70% water change daily in an environmental chamber at a temperature of 10±0.5°C. Exposure water was sampled at a time zero and time 24h (following a water change and before the subsequent water change), over four separate time points throughout the exposure period, and frozen at –20 °C until analysis. Water quality measurements were taken weekly. As alevins approached swim-up, they were fed *Artemia nauplii* once per day, then increased to twice per day after one week. Upon the completion of the exposure, fry were euthanized in 150 mg/L buffered tricaine mesylate (MS-222), weighed, and measured for total and standard length.

### Juvenile Acute Exposure

Following the sub-chronic study, lake trout from the same genetic pool were exposed to 6PPD-q for 96 hours at an age of eight weeks post-hatch. Based on results from the sub-chronic study, fish were exposed to nominal 6PPD-q concentrations of 0.1, 0.3, 0.9, and 2.7 μg/L. Tanks were again maintained in an environmental chamber at a temperature of 10±0.5°C. under semi-static conditions, with a daily water change of 70%. Fish were not fed over the exposure period.

Following initiation of the exposure, fry were monitored for behavioral changes and mortality hourly for the first seven hours, and twice daily for the remaining 96 hours. Exposure water was again sampled at a time zero and time 24h, over three time points throughout the exposure period, and frozen at –20 °C. Upon the completion of the exposure, surviving fry were euthanized in 150 mg/L of buffered MS-222, weighed, and measured for total and standard length.

### Chemical Analysis

Water samples, consisting of 950 μL exposure water, and 50 μL of deuterium-labeled internal standard solution (1mg/L) were frozen at -20 °C in amber autosampler vials. Analytical confirmation of exposure concentrations was achieved using ultra-high-performance liquid chromatography in tandem with high-resolution mass spectrometry following the methods described by ^5^. Calibration standard concentrations were within 15% of nominal, and target quantification of 6PPD-q was done using TraceFinder 4.1. Concentrations were reported as time-weighted average (TWA) across all replicates in a treatment.

### Data Analysis and Statistics

Chronic percent mortality was calculated using the Kaplan-Meier function S_t+1_ = St*((N_t+1_-D_t+1_)/N_t+1_), and LC50s were interpolated using the [agonist] vs. response – variable slope (four parameters) model via Prism 10.1.2 for Windows (GraphPad Software, Boston, MA).

## RESULTS

### Exposure Concentration Validation

LC-MS/MS quantification resulted in TWAs of 0.22, 0.58, 1.3, 3.4, 7.6, and 13.5 μg/L 6PPD-q for the sub-chronic exposure. For the 96-hour juvenile exposure, TWA concentrations of 0.05 μg/L in the lowest treatment, as levels were below the limit of detection, this represents half the limit of detection, and 0.16, 0.55, and 2.1 μg/L in the three highest exposure groups. Concentrations in the lowest treatment were below the limit of detection, and were reported as half the limit of detection (0.05 μg/L). Water quality parameters were ammonia 0.03±0.18 ppm, nitrite 0.47±0.58 ppm, nitrate 0.73±0.23 ppm, total hardness 92.67±9.27, pH 8.23±0.20, and DO 93±4.65%.

### Sub-chronic toxicity

Lake trout were sensitive to exposure with 6PPD-q, resulting in significant mortality and deformities (Figures 1-3). Significant mortality was observed four days post-initiation of exposure at hatch and continued throughout the exposure period with a 45-day LC_50_ of 0.39 μg/L [95% Confidence Interval (CI) of 0.26 to 0.41 μg/L] (Figure 2). No behavioral changes were noted. Several types of deformities were observed, with yolk sac edema, spinal curvature, and pooling of blood in the tail and eye being the most prominent (Figure 3). These deformities were noted within 72-hours of commencing exposure, and continued to occur until day 28, after which no new deformities were observed. The most common deformity was yolk sac edema, which occurred in all but one treatment. After 45 days of exposure, there was no significant difference in total length or weight among treatments in those fish remaining.

**Figure 1.**
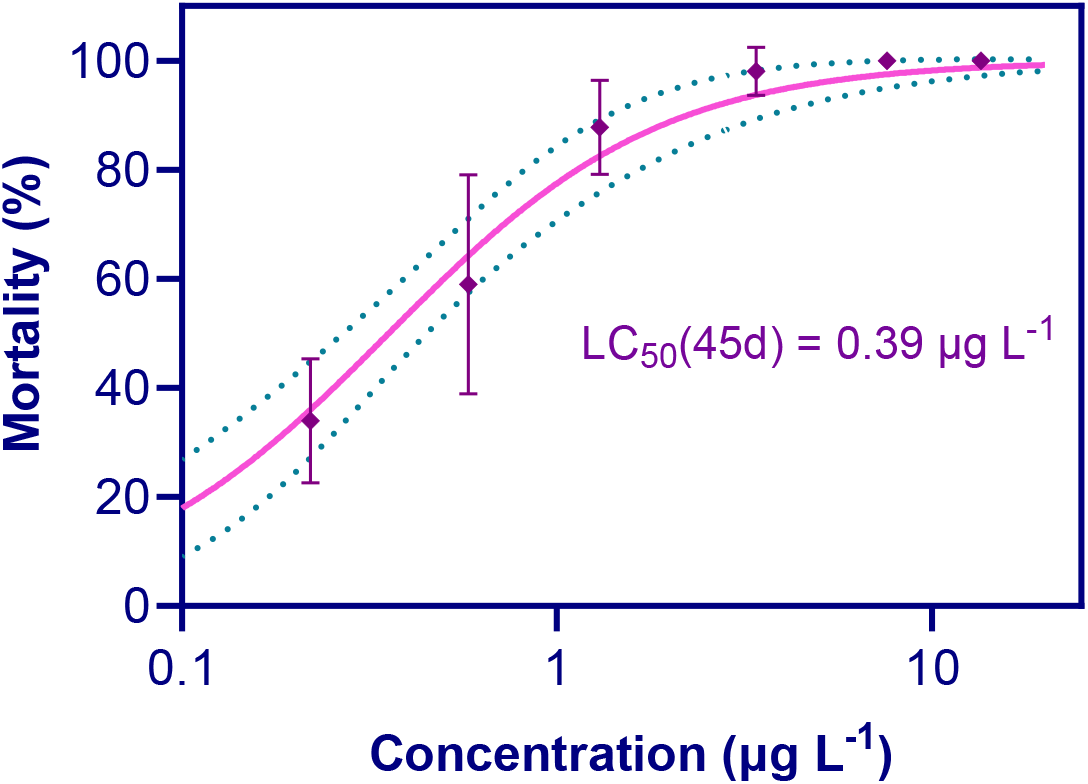
Percent mortality of early-life stage lake trout exposed to 6PPD-q for 45 days beginning at hatch. The 45-day LC_50_ was 0.30 μg/L, with a 95% CI of 0.30 to 0.48 μg/L. Points represent mean of replicates for each concentration; standard deviation is represented by the bars. The dotted lines indicate 95% confidence interval.

**Figure 2.**
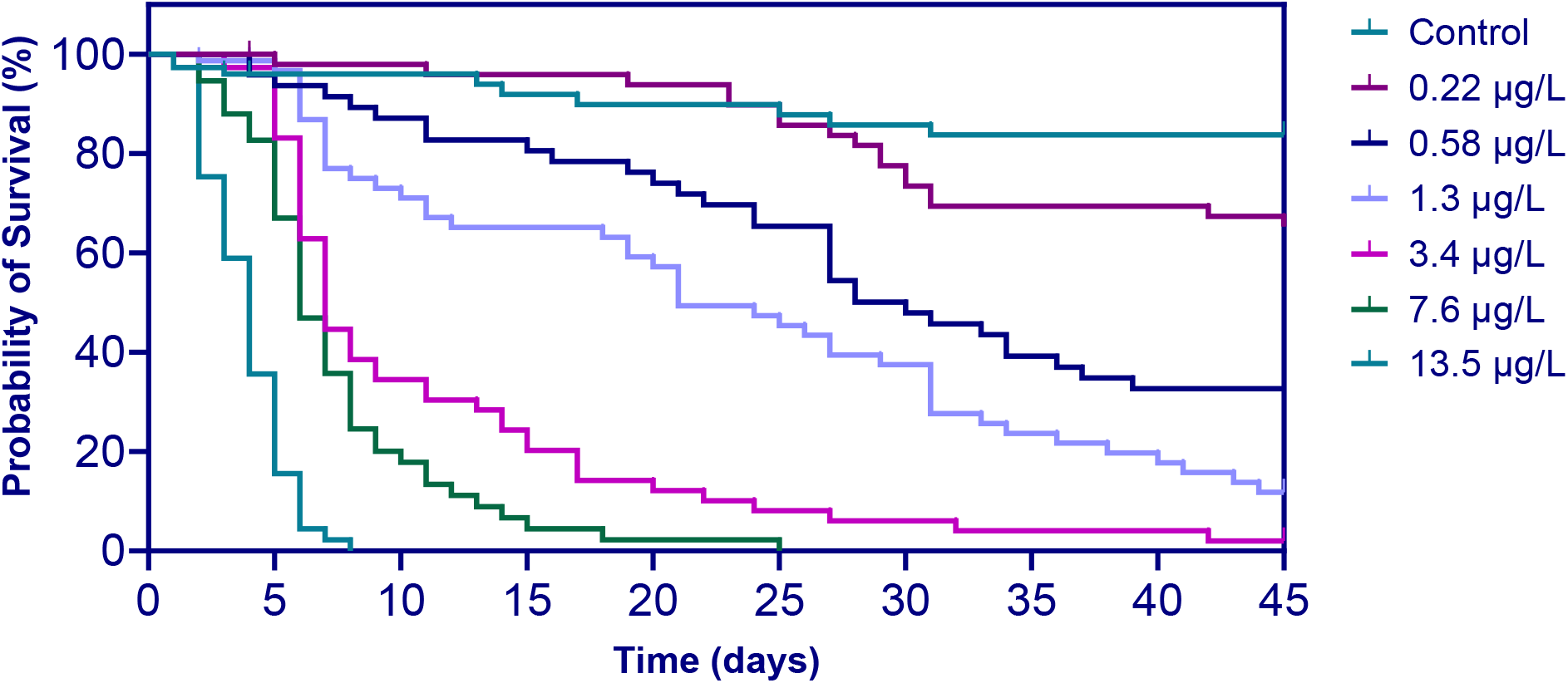
Kaplan-Meier survival analysis for early-life stag lake trout exposed to 6PPD-q for 45 days beginning at hatch.

**Figure 3.**
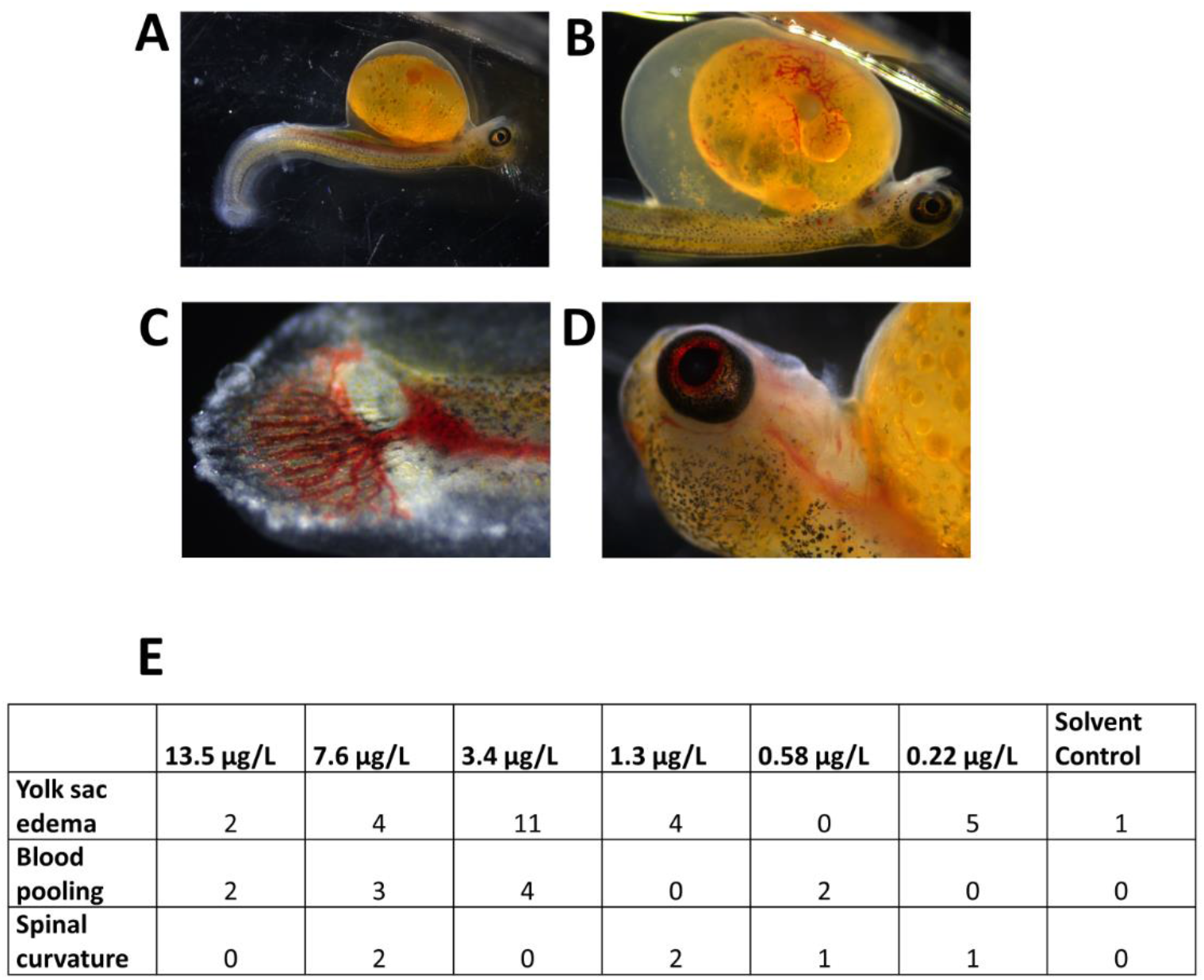
Examples of morphological changes observed in early-life stage lake trout exposed to 6PPD-q for 45 days beginning at hatch. **A**. Example of spinal curvature (7.6 μg/L treatment). **B**. Example of yolk sac edema (3.4 μg/L treatment). **C**. Example of caudal fin pathology (3.4 μg/L treatment). **D**. Pooling of blood in the eye (7.6 μg/L treatment). **E**. Total incidences of morphological abnormalities over 45 days in each treatment (note: these represent total numbers and were not normalized to mortalities).

### Juvenile Acute Exposure

Exposure to 6PPD-q in the juvenile life stage resulted in a concentration-dependent increase in mortality. Acute toxicity of 6PPD-q was apparent after only one hour of exposure of juvenile lake trout exposed to a concentration of 2.1 μg/L (Figure 4). The two highest treatment concentrations of 2.1 and 0.55 μg/L resulted in behavioural symptoms consistent with other 6PPD-q acute exposures, including loss of coordination, gasping, and surface swimming. All fish which exhibited these symptoms died within six hours, and no symptomatic fish recovered. No fish in the control, 0.16, or 0.05 μg/L treatments exhibited any symptoms or changes in behavior. There was no significant difference in total length or weight across treatments following completion of the exposure (data not shown). The 96-hour LC_50_ for the 8-week-old juvenile lake trout was 0.50 μg/L (95% CI of 0.42 to 0.60 μg/L) (Figure 5).

**Figure 4.**
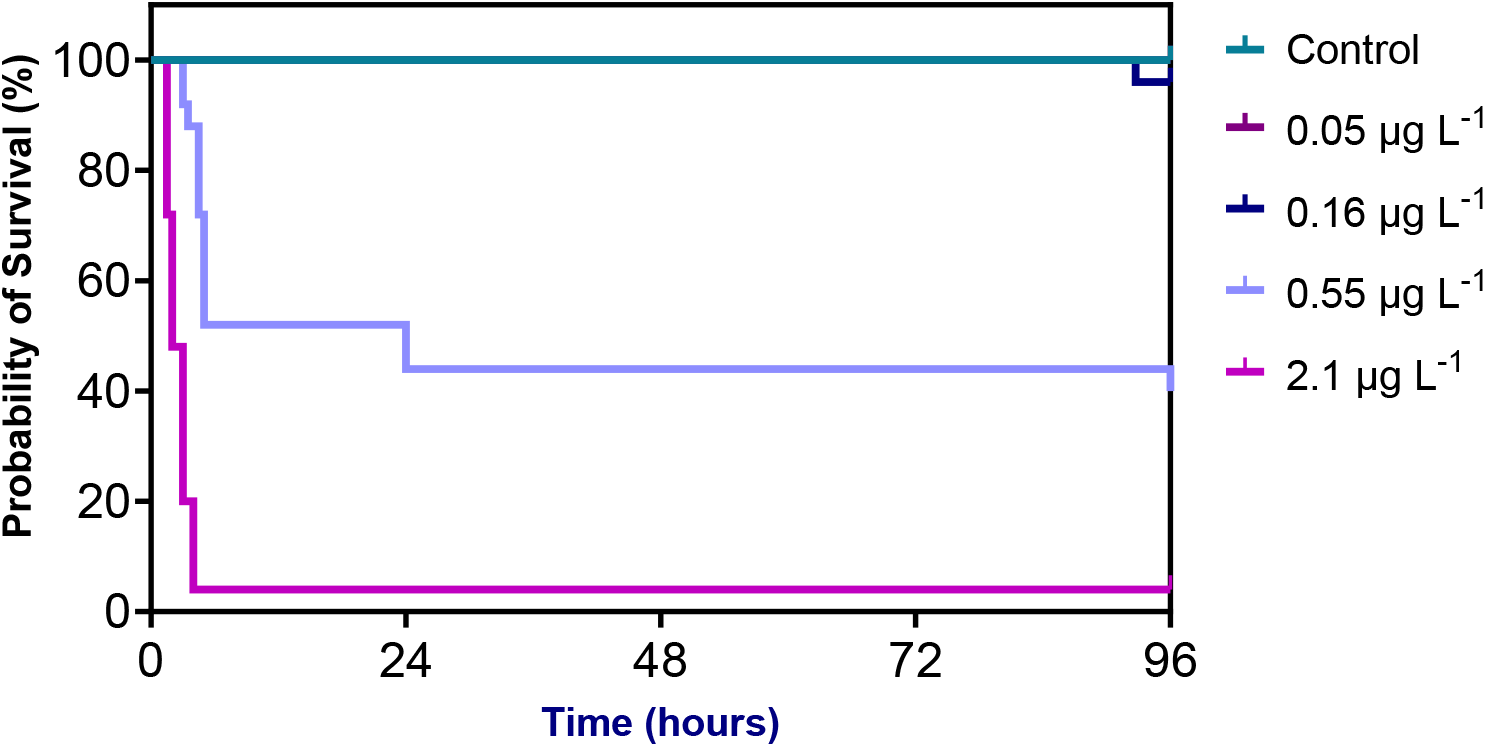
Time to survival analysis for 8-week-old lake trout exposed to 6PPD-q for 96 hours.

**Figure 5.**
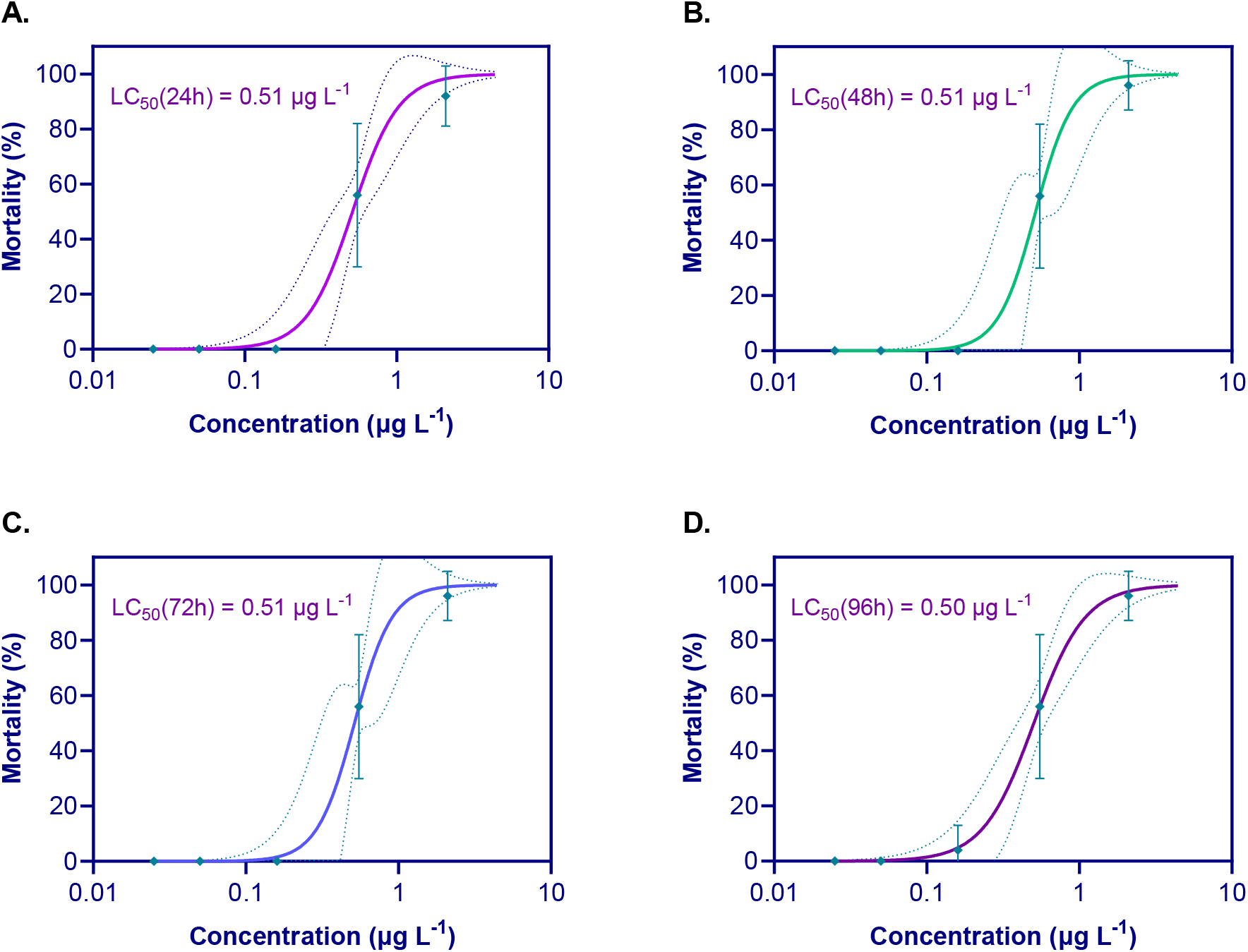
Percent mortality of 8-week-old lake trout exposed to 6PPD-q at times **A**. 24 hours **B**. 48 hours **C**. 72 hours **D**. 96 hours. Points represent mean of replicates for each concentration, standard deviation is represented by the bars. The dotted lines indicate 95% confidence interval.

## DISCUSSION AND CONCLUSIONS

The present study demonstrated that early-life stages of lake trout are sensitive to the exposure with low, environmentally relevant concentrations of 6PPD-q, adding another species to the list of salmonids susceptible to this emerging contaminant. Thus far, studies have elucidated a notable range of sensitive species within the *Oncorhynchus* genus, including coho salmon and rainbow trout ^8,15^, while other species in this genus such as Atlantic salmon (*Oncorhynchus keta*) and westslope cutthroat trout are not ^16,17^. Interestingly, char species also exhibit either tolerance, such as Arctic char ^8^ and southern Asian dolly varden ^9^, or high sensitivity, such as white-spotted char ^9^ and brook trout ^8^. The acute 24-96-hr LC_50_s of juvenile lake trout were very similar to the 24-hr LC_50_s of white-spotted char (0.51 μg/L) and brook trout (0.59 μg/L) ^8,9^, suggesting that these species of char may share some key similarity in terms of 6PPD-q toxicity. The wide variation among species is a consistent finding in the study of 6PPD-q as an aquatic contaminant, and so far, phylogeny alone does not seem to be a reliable predictor of sensitivity. Mitochondrial dysfunction has been proposed as a mechanism of action for 6PPD-q toxicity ^18^, which has been extrapolated upon ^10,19^ to include subsequent production of reactive oxygen species and changes in membrane permeability as a mechanism of toxicity. Given the conservation of mitochondrial and electron transport chain physiology among species, it is possible that toxicokinetic factors are a key aspect of the wide-ranging sensitivity of species. The ability to clear 6PPD-q at an adequate rate, or the production of a toxic metabolite(s) ^9,20^ may be the causative factor in determining sensitivity of a species.

While early-life stages cannot be directly compared to adults, juvenile lake trout were among the most sensitive species when comparing their acute LC_50_ with those obtained from previously reported experiments with adult or sub-adult fishes, surpassed only by juvenile and adult coho salmon ^11,15^. This high acute lethality may have significant implications for lake trout populations, especially in Canada. According to the Lake Ontario Technical Committee: Lake Trout Working Group, the failure of early-life survival is “likely the largest impediment to lake trout restoration in contemporary Lake Ontario,” ^21^. With both early-life stage sub-chronic and juvenile acute LC_50_ falling within environmentally relevant levels, exposure to 6PPD-q may further stress an already struggling population. The loss of early-life stage alevins not only hampers all subsequent adult population maintenance efforts through a reduction in recruitment but also severely diminishes population diversity by creating genetic bottlenecks ^22–24^.

In addition, it is possible that early-life stages of salmonids are exposed to greater concentrations of 6PPD-q compared to those that are measured in the water column, or over longer periods of time, given that TWP settle into sediments (Li et al., 2023). As TWP are washed into aquatic environments, the 6PPD on the surface of the TWP is converted to 6PPD-q, the more stable compound (Hu et al., 2023), which is still short-lived in the water column. This results in surface waters experiencing intermittent pulses of 6PPD-q, correlated with precipitation (Greer et al., 2023). However, the settling of TWPs in sediment may provide another, more constant and long-term source of 6PPD-q. It is worth noting that early-life stage lake trout alevins inhabit this gravel environment, in contrast to their adult counterparts which are predominantly found in the water column in deeper parts of lakes. The newly-laid eggs incubate in the gravel for several months, and after hatching, the alevins spend several weeks consuming their yolk-sac in the gravel before swimming up to feed exogenously. Therefore, the period from hatching to swim-up may be a critical time during which lake trout alevins are most vulnerable to the toxic effects of 6PPD-q.

While we observed significant mortality in both the juvenile acute and sub-chronic groups, only those alevins exposed from hatch, in a sub-chronic manner, exhibited significant pathologies, including spinal curvature, yolk sac edema, and pooling of blood in the caudal fin and eye. This may be due to both the life stage and the duration of exposure. We found that the earliest pathologies occurred within 72 hours of exposure in the post-hatch alevins, but not in juveniles. This suggests that early growth and development are significantly disrupted by 6PPD-q at this life stage. However, we also noted new pathologies up to 28 days into the exposure, indicating that chronic exposure to 6PPD-q can result in sub-lethal effects in lake trout fry. These sub-lethal effects could have population-level implications for overall survival in fry that are exposed to low, sub-chronic, or chronic levels of 6PPD-q. It is important to note that spinal curvature can impact a fish’s ability to swim effectively ^25^, affecting their ability to find food, escape from predators, and reproduce. This has broad implications for the long-term survival and development of lake trout fry. While none of these pathologies have been reported in associated with 6PPD-q exposure, pooling of blood in the caudal fin and eye are consistent with previous studies that have shown changes in vascular and blood-brain barrier permeability in coho salmon following exposure to 6PPD-q ^10,19^. It is worth noting that none of the fish exhibiting these pathologies survived the exposure, suggesting that the physiological changes which lead to the development of these pathologies, might be predictive, if not causative, of mortality.

In all previous studies, acute exposure of sensitive salmonid species to 6PPD-q resulted in similar symptomologies of gaping, loss of coordination, and surface swimming. We observed these same changes in our juvenile acute study, but no symptoms were observed in the sub-chronic exposure. In addition, we also did not observe mortality at the same rate in the newly hatched larvae versus the juveniles. This may be due to the under-development of the fish’s brain and nervous system or possibly an initial adaptive period in which 6PPD-q may be sequestered in lipid-rich tissues like the yolk sac. However, it is important to consider that deformities in the caudal fin, as well as pooling of blood in the eye, have the potential to impact a fish’s swimming ability and vision, respectively. Nevertheless, these changes only become relevant if any fry with these deformities are able to survive for a significant period of time.

### Future Outlook

This research demonstrated the high sensitivity of another salmonid species to the exposure with 6PPD-q, deepening the question as to what physiological components are involved in toxicity. Given the conservation of mitochondrial function and ROS effects in fishes, this wide range of species sensitivity may indicate a toxicokinetic driver of toxicity. Proposed by Hiki et al., the presence of a monohydroxylated metabolite in exposed fishes may indicate that toxicity stems from the production of a toxic metabolite in those fish species which are sensitive, and not in those tolerant ^9^. However, given the insensitivity of RT-Hep in contrast with RT-W1 gill, it is also possible that sensitivity is determined by an organism’s ability to clear 6PPD-q itself. The difference in sensitive and tolerant species may come down to something as simple as a change in a CYP isomer. This also brings into question potential differences in sensitivity between early life stages and adult, as if toxicity is driven by toxicokinetic factors, those early-life stages with reduced capacity for metabolism may also be an important factor in population-level effects of 6PPD-q exposure.

Thus, further studies looking at metabolite production in tolerant and sensitive species is key to determining the role of toxicokinetics in this unusual species sensitivity distribution. In addition, work to determine changes in gene expression could also help point toward a greater understanding of the mechanism(s) of toxicity. Differential gene expression could provide key insight as to these differences in sensitivity, as well as physiological changes, particularly those at a sub-lethal level. Further to this, there are a number of other salmonid species for which there is no sub-chronic or early-life stage information in regards to 6PPD-q exposure. Given that early-life stages fishes are, on average 60%, more sensitive to contaminants ^12^, the degree of risk to those species where the adults are known to be sensitive may be even greater. Understanding the potential risk to these populations is integral to protecting these species, and in research and recommendations for mitigation efforts toward limiting 6PPD-q exposure and Urban Runoff Mortality Syndrome.

## Acknowledgments

This project was supported partially by a financial contribution from Fisheries and Oceans Canada. Additional funding was provided to M.B. and M.H. through the Discovery Grants program of the Natural Sciences and Engineering Research Council of Canada (NSERC). M.B. is currently a faculty member of the Global Water Futures (GWF) program, which received funds from the Canada First Research Excellence Funds (CFREF). M.H. was supported by the Canada Research Chairs Program. The authors acknowledge the support of Zoë Henrikson, Dale Jefferson, and Azadeh Hatef (ATRF) in animal care. The authors also acknowledge the support of Government of Saskatchewan Fisheries Unit for assistance in procurement of embryos, as well as general lab support provided by Matthew Schultz, Julie Bowen, Kerstin Bluhm, Phillip Ankley, and Katherine Raes.

## References

(1) Feist, B. E.; Buhle, E. R.; Arnold, P.; Davis, J. W.; Scholz, N. L. Landscape Ecotoxicology of Coho Salmon Spawner Mortality in Urban Streams. PLoS One 2011, 6 (8), 23424. 10.1371/journal.pone.0023424.

(2) Feist, B. E.; Buhle, E. R.; Baldwin, D. H.; Spromberg, J. A.; Damm, S. E.; Davis, J. W.; Scholz, N. L. Roads to Ruin: Conservation Threats to a Sentinel Species across an Urban Gradient. Ecol. Appl. 2017, 27 (8), 2382–2396. 10.1002/EAP.1615.

(3) Scholz, N. L.; Myers, M. S.; McCarthy, S. G.; Labenia, J. S.; McIntyre, J. K.; Ylitalo, G. M.; Rhodes, L. D.; Laetz, C. A.; Stehr, C. M.; French, B. L.; McMillan, B.; Wilson, D.; Reed, L.; Lynch, K. D.; Damm, S.; Davis, J. W.; Collier, T. K. Recurrent Die-Offs of Adult Coho Salmon Returning to Spawn in Puget Sound Lowland Urban Streams. PLoS One 2011, 6 (12), 28013. 10.1371/journal.pone.0028013.

(4) Tian, Z.; Zhao, H.; Peter, K. T.; Gonzalez, M.; Wetzel, J.; Wu, C.; Hu, X.; Prat, J.; Mudrock, E.; Hettinger, R.; Cortina, A. E.; Biswas, R. G.; Kock, F. V. C.; Soong, R.; Jenne, A.; Du, B.; Hou, F.; He, H.; Lundeen, R.; Gilbreath, A.; Sutton, R.; Scholz, N. L.; Davis, J. W.; Dodd, M. C.; Simpson, A.; Mcintyre, J. K.; Kolodziej, E. P. A Ubiquitous Tire Rubber-Derived Chemical Induces Acute Mortality in Coho Salmon. Science (80-.). 2021, 375 (6582), eabo5785. 10.1126/science.abo5785.

(5) Challis, J. K.; Popick, H.; Prajapati, S.; Harder, P.; Giesy, J. P.; McPhedran, K.; Brinkmann, M. Occurrences of Tire Rubber-Derived Contaminants in Cold-Climate Urban Runoff. Environ. Sci. Technol. Lett. 2021, 8 (11), 961–967. 10.1021/acs.estlett.1c00682.

(6) Hu, X.; Zhao, H. N.; Tian, Z.; Peter, K. T.; Dodd, M. C.; Kolodziej, E. P. Transformation Product Formation upon Heterogeneous Ozonation of the Tire Rubber Antioxidant 6PPD (N-(1,3-Dimethylbutyl)-N′-Phenyl-p-Phenylenediamine). Environ. Sci. Technol. Lett. 2022. 10.1021/acs.estlett.2c00187.

(7) Johannessen, C.; Helm, P.; Lashuk, · Brent; Yargeau V.; Metcalfe, C. D. The Tire Wear Compounds 6PPD-Quinone and 1,3-Diphenylguanidine in an Urban Watershed. Arch. Environ. Contam. Toxicol. 2021, 1, 3. 10.1007/s00244-021-00878-4.

(8) Brinkmann, M.; Montgomery, D.; Selinger, S.; Miller, J. G. P.; Stock, E.; Alcaraz, A. J.; Challis, J. K.; Weber, L.; Janz, D.; Hecker, M.; Wiseman, S. Acute Toxicity of the Tire Rubber-Derived Chemical 6PPD-Quinone to Four Fishes of Commercial, Cultural, and Ecological Importance. Environ. Sci. Technol. Lett. 2022, 9 (4), 333–338. 10.1021/acs.estlett.2c00050.

(9) Hiki, K.; Yamamoto, H. The Tire-Derived Chemical 6PPD-Quinone Is Lethally Toxic to the White-Spotted Char Salvelinus Leucomaenis Pluvius but Not to Two Other Salmonid Species. Environ. Sci. Technol. Lett. 2022, 9 (12), 1050–1055. 10.1021/acs.estlett.2c00683.

(10) Greer, J. B.; Dalsky, E. M.; Lane, R. F.; Hansen, J. D. Tire-Derived Transformation Product 6PPD-Quinone Induces Mortality and Transcriptionally Disrupts Vascular Permeability Pathways in Developing Coho Salmon. Environ. Sci. Technol. 2023, 57 (30), 10940–10950. 10.1021/acs.est.3c01040.

(11) Lo, B. P.; Marlatt, V. L.; Liao, X.; Reger, S.; Gallilee, C.; Ross, A. R. S.; Brown, T. M. Acute Toxicity of 6PPD-Quinone to Early Life Stage Juvenile Chinook (Oncorhynchus Tshawytscha) and Coho (Oncorhynchus Kisutch) Salmon. Environ. Toxicol. Chem. 2023, 42 (4), 815–822. 10.1002/etc.5568.

(12) Hutchinson, T. H.; Solbé, J.; Kloepper-Sams, P. J. Analysis of the ECETOC Aquatic Toxicity (EAT) Database. III - Comparative Toxicity of Chemical Substances to Different Life Stages of Aquatic Organisms. Chemosphere 1998, 36 (1), 129–142. 10.1016/S0045-6535(97)10025-X.

(13) Fisheries Canada. Lake Trout. https://www.dfo-mpo.gc.ca/species-especes/profiles-profils/lake-trout-touladi-eng.html (accessed 2024-01-16).

(14) Ontario Ministry of Natural Resources and Forestry. Lake Trout management strategy. https://www.ontario.ca/document/fisheries-management-plan-fisheries-management-zone-18/lake-trout-management-strategy#section-4 (accessed 2023-10-03).

(15) Tian, Z.; Gonzalez, M.; Rideout, C. A.; Zhao, H. N.; Hu, X.; Wetzel, J.; Mudrock, E.; James, C. A.; McIntyre, J. K.; Kolodziej, E. P. 6PPD-Quinone: Revised Toxicity Assessment and Quantification with a Commercial Standard. Environ. Sci. Technol. Lett. 2022. 10.1021/acs.estlett.1c00910.

(16) Montgomery, D.; Ji, X.; Cantin, J.; Philibert, D.; Foster, G.; Selinger, S.; Jain, N.; Miller, J.; McIntyre, J.; de Jourdan, B.; Wiseman, S.; Hecker, M.; Brinkmann, M. Interspecies Differences in 6PPD-Quinone Toxicity Across Seven Fish Species: Metabolite Identification and Semiquantification. Environ. Sci. Technol. 2023. 10.1021/acs.est.3c06891.

(17) Foldvik, A.; Kryuchkov, F.; Sandodden, R.; Uhlig, S. Acute Toxicity Testing of the Tire Rubber–Derived Chemical 6PPD-Quinone on Atlantic Salmon (Salmo Salar) and Brown Trout (Salmo Trutta). Environ. Toxicol. Chem. 2022, 41 (12), 3041–3045. 10.1002/etc.5487.

(18) Mahoney, H.; Da Silva Junior, F. C.; Roberts, C.; Schultz, M.; Ji, X.; Alcaraz, A. J.; Montgomery, D.; Selinger, S.; Challis, J. K.; Giesy, J. P.; Weber, L.; Janz, D.; Wiseman, S.; Hecker, M.; Brinkmann, M. Exposure to the Tire Rubber-Derived Contaminant 6PPD-Quinone Causes Mitochondrial Dysfunction in Vitro. Environ. Sci. Technol. Lett. 2022, 9 (9), 765–771. 10.1021/acs.estlett.2c00431.

(19) Blair, S. I.; Barlow, C. H.; McIntyre, J. K. Acute Cerebrovascular Effects in Juvenile Coho Salmon Exposed to Roadway Runoff. Can. J. Fish. Aquat. Sci. 2021, 78 (2), 103–109. 10.1139/cjfas-2020-0240.

(20) Nair, P.; Sun, J.; Xie, L.; Kennedy, L.; Kozakiewicz, D.; Kleywegt, S.; Hao, C.; Byun, H.; Barrett, H.; Baker, J.; Monaghan, J.; Krogh, E.; Song, D.; Peng, H. Synthesis and Toxicity Evaluation of Tire Rubber-Derived Quinones. ChemRxiv 2023, 1–28.

(21) Lake Ontario Technical Committee: Lake Trout Working Group. Research Priorities for Lake Trout Restoration in Lake Ontario, 2022; 2022. http://www.glfc.org/pubs/lake_committees/ontario/2022_LOTC_Research_Priorities.pdf (accessed 2023-04-16).

(22) Petit-Marty, N.; Liu, M.; Tan, I. Z.; Chung, A.; Terrasa, B.; Guijarro, B.; Ordines, F.; Ramírez-Amaro, S.; Massutí, E.; Schunter, C. Declining Population Sizes and Loss of Genetic Diversity in Commercial Fishes: A Simple Method for a First Diagnostic. Front. Mar. Sci. 2022, 9 (May), 1–11. 10.3389/fmars.2022.872537.

(23) McMillan, A. M.; Bagley, M. J.; Jackson, S. A.; Nacci, D. E. Genetic Diversity and Structure of an Estuarine Fish (Fundulus Heteroclitus) Indigenous to Sites Associated with a Highly Contaminated Urban Harbor. Ecotoxicology 2006, 15 (6), 539–548. 10.1007/s10646-006-0090-4.

(24) Nacci, D.; Coiro, L.; Champlin, D.; Jayaraman, S.; McKinney, R.; Gleason, T. R.; Munns, W. R.; Specker, J. L.; Cooper, K. R. Adaptations of Wild Populations of the Estuarine Fish Fundulus Heteroclitus to Persistent Environmental Contaminants. Mar. Biol. 1999, 134 (1), 9–17. 10.1007/s002270050520.

(25) Toften, H.; Jobling, M. Development of Spinal Deformities in Atlantic Salmon and Arctic Charr Fed Diets Supplemented with Oxytetracycline. J. Fish Biol. 1996, 49 (4), 668–677. 10.1006/jfbi.1996.0195.

